# The Dynamics of Error Processing in the Human Brain as Reflected by High-Gamma Activity in Noninvasive and Intracranial EEG

**DOI:** 10.1101/166280

**Authors:** Martin Völker, Lukas D. J. Fiederer, Sofie Berberich, Jiří Hammer, Joos Behncke, Pavel Kršek, Martin Tomášek, Petr Marusič, Peter C. Reinacher, Volker A. Coenen, Moritz Helias, Andreas Schulze-Bonhage, Wolfram Burgard, Tonio Ball

**Author notes:** These authors contributed equally to this work (Lukas D. J. Fiederer, Sofie Berberich, Jiří Hammer). Corresponding author: Martin Völker, Translational Neurotechnology Lab, Engelbergerstr. 21, 79106 Freiburg, Germany.

## Abstract

Error detection in motor behavior is a fundamental cognitive function heavily relying on cortical information processing. Neural activity in the high-gamma frequency band (HGB) closely reflects such local cortical processing, but little is known about its role in error processing, particularly in the healthy human brain. Here we characterize the error-related response of the human brain based on data obtained with noninvasive EEG optimized for HGB mapping in 31 healthy subjects (15 females, 16 males), and additional intracranial EEG data from 9 epilepsy patients (4 females, 5 males). Our findings reveal a comprehensive picture of the global and local dynamics of error-related HGB activity in the human brain. On the global level as reflected in the noninvasive EEG, the error-related response started with an early component dominated by anterior brain regions, followed by a shift to parietal regions, and a subsequent phase characterized by sustained parietal HGB activity. This phase lasted for more than 1 s after the error onset. On the local level reflected in the intracranial EEG, a cascade of both transient and sustained error-related responses involved an even more extended network, spanning beyond frontal and parietal regions to the insula and the hippocampus. HGB mapping appeared especially well suited to investigate late, sustained components of the error response, possibly linked to downstream functional stages such as error-related learning and behavioral adaptation. Our findings establish the basic spatio-temporal properties of HGB activity as a neural correlate of error processing, complementing traditional error-related potential studies.

**Significance Statement:** There is great interest to understand how the human brain reacts to errors in goal-directed behavior. An important index of cortical and subcortical information processing is fast oscillatory brain activity, particularly in the high-gamma band (above 50 Hz). Here we show that it is possible to detect signatures of errors in event-related high-gamma responses with noninvasive techniques, characterize these responses comprehensively, and validate the EEG procedure for the detection of such signals. In addition, we demonstrate the added value of intracranial recordings pinpointing the fine-grained spatio-temporal patterns in error-related brain networks. We anticipate that the optimized noninvasive EEG techniques as described here will be helpful in many areas of cognitive neuroscience where fast oscillatory brain activity is of interest.

## Introduction

Error processing is a fundamental brain function. A breakthrough in research on error processing in the human brain was the independent discovery of the “error-related negativity” (ERN) (Gehring et al., 1993), or error negativity (Ne) (Falkenstein et al., 1991) in noninvasive electroencephalography (EEG). The ERN/Ne is a negative deflection above the fronto-central midline, peaking shortly after the electromyogram (EMG) onset of an erroneous response, followed by the error positivity (Pe) (Falkenstein et al., 1991) with parietal maximum. ERN/Ne and Pe are often assumed to reflect sequential functional aspects of error processing, including precursors of error detection such as conflict monitoring and explicit error detection itself. In contrast to the ERN/Ne, the Pe was linked to conscious error processing (Nieuwenhuis et al., 2001). Moreover, the Pe might reflect evidence strength during error detection and could thus provide input to further downstream stages, e.g., to the evaluation of the significance of errors and the implementation of behavioral reactions (Steinhauser and Yeung, 2010).

To further dissect error-related processing both in healthy subjects and in patients with a broad spectrum of brain disorders (Alain et al., 2002; Hajcak et al., 2003; Shiels and Hawk, 2010), subsequent studies increasingly utilized time-frequency decomposition of error-related EEG responses. These studies revealed spatial and dynamical behavior of lower frequency bands like delta, theta, alpha and beta (Table 2) unfolding alongside the time-domain (ERN/Ne & Pe) evoked electrical field potential changes. There is only limited data on the role of higher frequencies in error processing, coming from intracranial recordings in neurological patients (Milekovic et al., 2013; Bastin et al., 2017) and we are not aware of any other study using noninvasive EEG recorded from the healthy human brain.

**Table 2:**
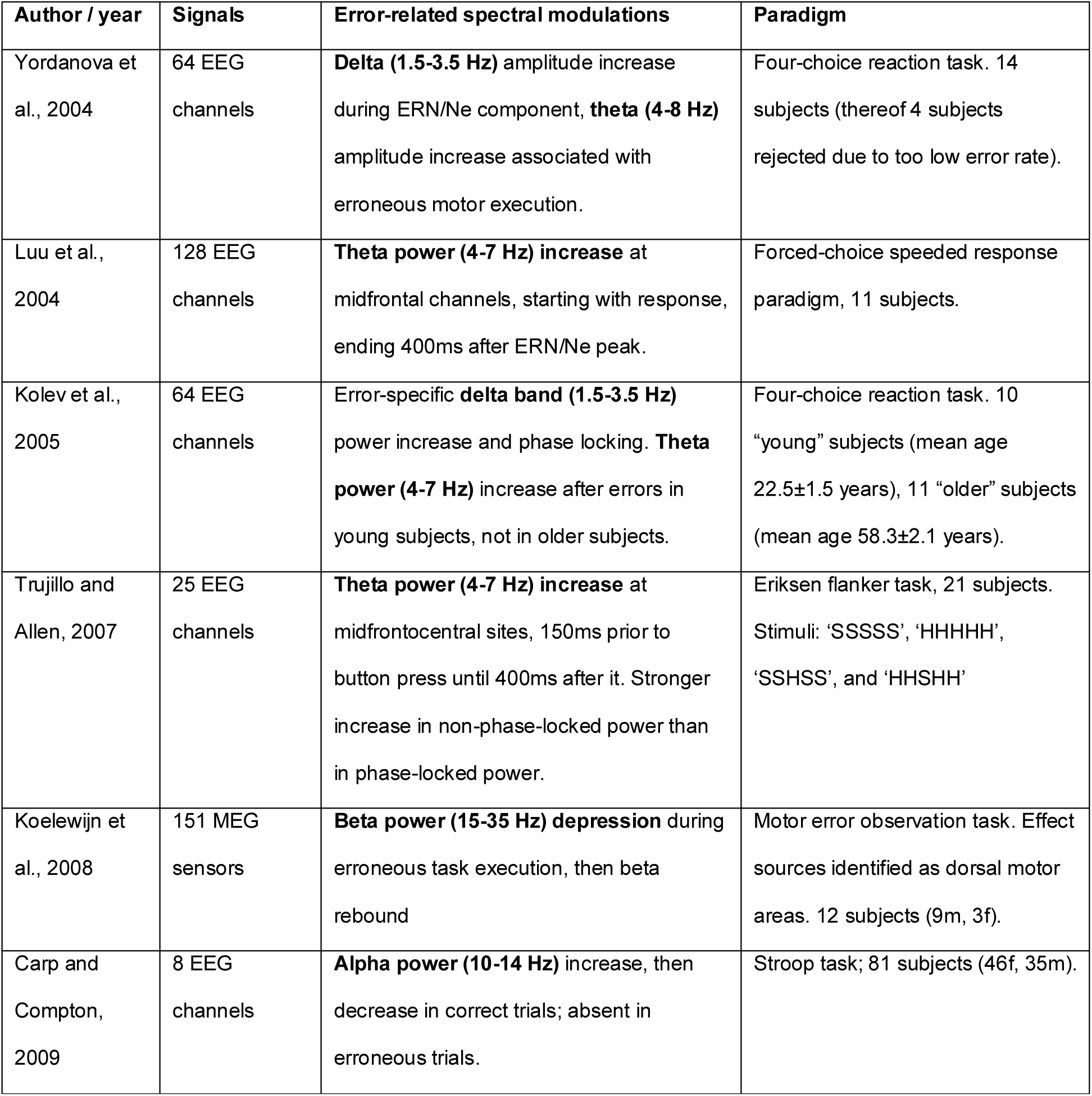
**Excerpt of studies about error-related spectral power modulations in EEG & MEG.**
As frequency band definitions were not consistent amongst the publications, the respective definition is stated within parentheses.

However, gamma-band frequencies may be especially important to understand cortical function in general, including error processing. A large body of empirical evidence indicates that high-gamma band (HGB, including the 50-150 Hz range) activity is a spatially and temporally specific index of the underlying, functionally relevant neural networks (Crone et al., 1998, 2006; Brunel and Wang, 2003). Compared to lower frequencies, however, detecting HGB power modulations in noninvasive EEG is challenging for several reasons that are related to the more focal spatial distribution of cortical high-frequency sources (Crone et al., 2006), their much smaller power, (Freeman et al., 2000), and their greater susceptibility to artifacts, such as from muscle activity (Goncharova et al., 2003) or microsaccades. The latter particularly can mimic physiological responses within the HGB range (Yuval-Greenberg et al., 2008).

To overcome these problems, we carefully optimized the procedure of EEG acquisition and analysis for the detection of high-frequency EEG modulations, combining high-resolution EEG acquisition, optimized electromagnetic shielding, low-noise amplifier systems, as well as high-precision eye tracking simultaneously acquired to the EEG data to tightly control for ocular artifacts. Utilizing this optimized setup, we re-examined a classical paradigm to elicit error responses in a large group (n=35) of healthy subjects. Furthermore, to validate our noninvasive EEG findings, we also ran the same paradigm in patients with intracranially implanted electrodes.

Our findings clearly demonstrate that error-related HGB brain responses can be detected in noninvasive EEG recorded from healthy subjects; importantly, we rule out ocular including micro-saccadic effects as an explanation for the observed HGB responses, as well as corroborate our noninvasive observations by intracranial EEG data. For the first time, our findings reveal a clear picture of the global dynamics of the error-related HGB response of the human brain, starting from an early response dominated by anterior brain regions, over a shift to medial parietal regions parallel to the Pe, and finally to a subsequent phase characterized by sustained parietal HGB activity. This phase lasts for more than 1 s after the onset of the error event and constitutes a novel candidate signal of the downstream processes following the classical Pe. Combined investigation of both the classical error-related potentials and of HGB modulations thus promises to shed new light on error processing in the human brain.

## Materials & Methods

### Subjects

In the noninvasive EEG study, 35 healthy subjects participated; thereof, 4 subjects had to be excluded because of extensive muscular or ocular artifacts. Thus, data of 31 subjects (mean age 24.6 years, standard deviation (SD) = 3.1 years, 15 females) were further analyzed. Handedness was assessed according to a modified Edinburgh handedness questionnaire (Oldfield, 1971); 28 subjects were right-handed, 3 were lefthanded. All stated not to have neurological or psychiatric diseases and not to be under the influence of medication affecting the central nervous system.

In the intracranial EEG study, 9 right-handed patients (mean age 27.0 years, SD = 7.7 years, 4 females) with pharmacoresistant epilepsy were recruited. They were implanted with intracranial electrodes in Freiburg, Germany, or in Prague, Czech Republic.

All subjects and patients gave their written informed consent before participating in the study. The study was approved by the local ethics committees.

### Experimental design

To probe error-related processing, we used the Eriksen flanker task (Eriksen and Eriksen, 1979) (Fig. 1) as employed in a pioneering study in this field of research (Gehring et al., 1993) as well as in many follow-up publications (Kopp et al., 1996; Botvinick et al., 1999; Gehring and Knight, 2000; Nieuwenhuis et al., 2002; Ridderinkhof et al., 2002; Herrmann et al., 2004b; Albrecht et al., 2009; Maier et al., 2012; Zavala et al., 2013). At the beginning of each trial, a cue in the form of an asterisk was shown to the subjects in the center of a 19-inch monitor (4:3-screen ratio, 60-Hz frame rate) for 1 s. After that, one of four stimuli was shown for 100 ms, each with a probability of 0.25. Two of the stimuli were *congruent stimuli* (LLLLL & RRRRR) and the other two *incongruent stimuli* (RRLRR & LLRLL). To respond, subjects used their left or right index finger to press the left or right analog shoulder button on a wireless gamepad (Logitech F710, Apples, Switzerland) if the central letter of the stimulus was an “L” or “R”, respectively. The deflection threshold of the analog joystick button was set to 10% of its maximal deflection.

**Figure 1 :**
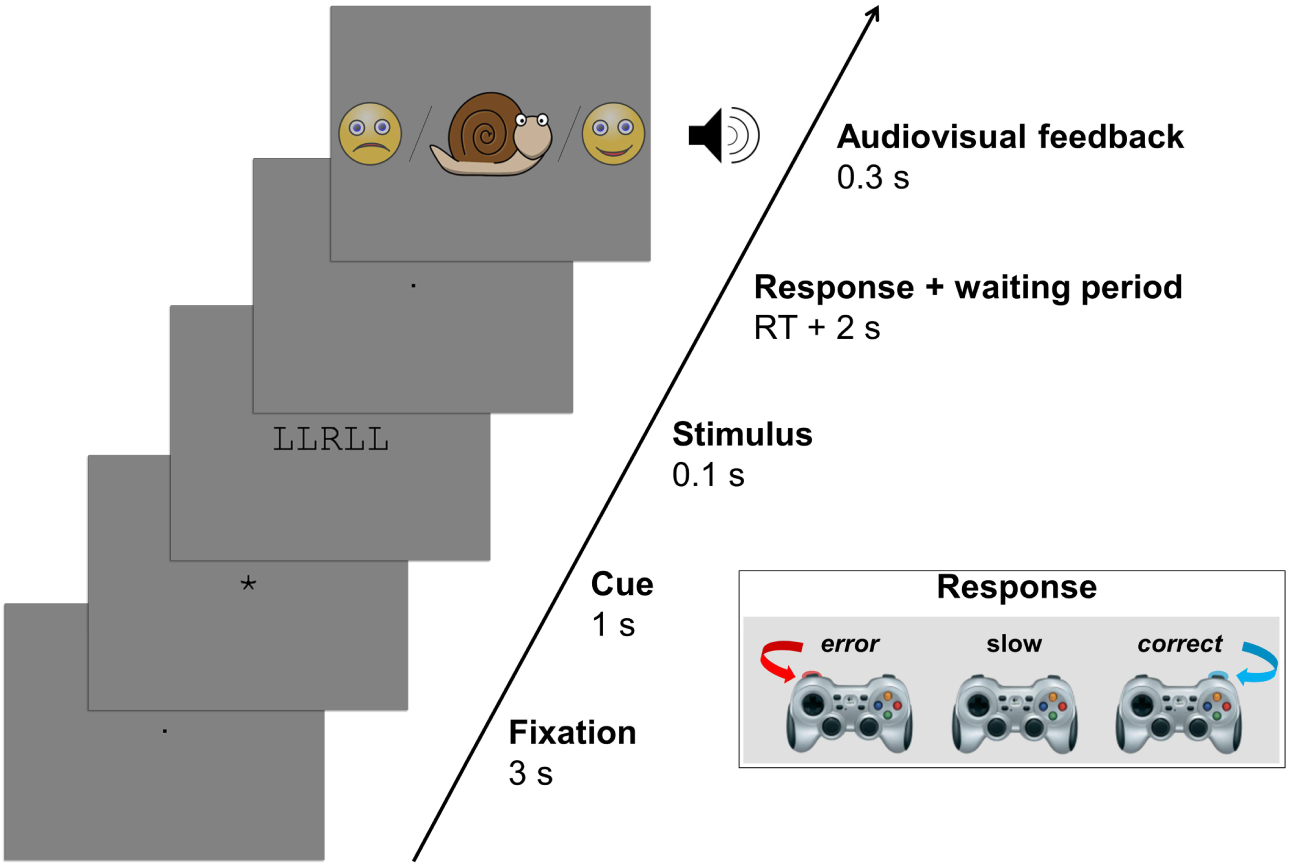
Flowchart of the Eriksen flanker task paradigm used to elicit errors. After a fixation period (3 s), and a cue period (1 s), the stimulus appeared for a short period of time (100 ms). Subjects had only a limited amount of time, individually set to their reaction time (RT) in a familiarization phase of the experiment, to press a gamepad button with their left index finger if the central letter of the stimulus was an “L” or with their right index finger if the central letter was an “R” (see example with ‘R’ target letter in inset). 2 s after the button press, one of the three types of audiovisual feedback indicated to the subjects whether their response was correct, incorrect, or too slow.

As proposed by (Herrmann et al., 2004b), we set an individual reaction time limit for each subject. This response time limit was determined as the mean response time in a 32-trial training session. Subjects were instructed to respond as fast and as accurately as possible. The subjects were also introduced to a scoring point system, and instructed to get as many points as possible. For each correct response, subjects gained 5 points, and lost 5 points in the event of an erroneous response. By missing the individual reaction time limit, subjects lost 10 points; this stronger penalty was introduced to keep the subject under time pressure, hence inducing errors. In each break after a recording run, the score and performance of the respective run was shown to the subjects along with a comparison to their total performance up to this run.

Two seconds after their response, the subjects received an audiovisual feedback according to their performance. If the response was fast enough and correct, the feedback consisted of a smiling face icon and a 1-kHz sine tone, the feedback for an erroneous but fast enough response consisted of a sad face and a 500-Hz sine tone. For responses slower than the individual reaction time limit, a cartoon of a snail was displayed accompanied by a 5-kHz sine tone. The two-second delay was introduced to avoid an influence of the feedback on error-related brain responses. We did not verify the error awareness prior to the feedback directly; however, all subjects reported that they were aware of most errors directly after the motor response.

Before the start of each trial, only the fixation dot was shown for 3 s. In the case of healthy subjects, one session consisted of 100 trials, after which the subjects had the possibility to have a break. Each experiment included 10 sessions, so that altogether 1000 trials were collected for each subject; on average, the error rate was 22.23 ± 0.11 (mean ± SD) %. Recording sessions with epilepsy patients were shorter and included overall fewer trials depending on the condition of the respective patient. On average, patients completed 369 ± 111 trials, thereof 218 ± 74 correct trials and 57 ± 32 error trials, with a total error rate of 20.95 ± 0.11 %.

### Recording and preprocessing of noninvasive EEG

The noninvasive EEG setup was optimized for the measurement of high-frequency responses. We used NeurOne amplifiers (Mega Electronics Ltd., Kuopio, Finland) with a 24-bit resolution and low input noise (root mean square < 0.6 μV between 0.16-200 Hz). We recorded 128 EEG channels with the waveguard EEG cap (ANT Neuro, Enschede, Netherlands) at a sampling rate of 5 kHz (AC, 1250-Hz anti-aliasing low-pass filter). The cap held sintered Ag/AgCl electrode elements and was available in three different sizes to be suitable for variable head sizes. Electrode position Cz was used as recording reference; the ground was located between AFz and Fz. Whenever possible, impedances were kept below 5 kΩ. Additional measurements included electrooculography (EOG) with 4 electrodes around the eyes, 2-channel electrocardiography (ECG) and bipolar electromyography (EMG) above the forearm flexor muscles of both arms and above the gastrocnemius muscle of both legs; EMG, EOG and ECG were recorded with self-adhesive electrodes.

The EEG recordings took place in an electromagnetically shielded cabin (“mrShield” -CFW Trading Ltd, Heiden, Switzerland) to reduce electromagnetic artifact contamination. All exchange of information between inside and outside of the cabin was done with fiber optic cables to sustain the shielding. Also, electrical devices inside the cabin, such as EEG amplifier, eye tracker and loudspeakers, were powered by DC batteries to prevent 50-Hz power line artifacts from interfering with the EEG signal. The cabin furthermore dampens sounds and vibrations to protect the subject from external noise.

Both control of the experiment as well as data analysis was carried out using Matlab R2014a (The MathWorks Inc., Natick, USA, RRID:SCR_001622). Implementation of the paradigm was done within the Psychophysics Toolbox (Brainard, 1997, RRID:SCR_002881). Synchronization of the EEG data and the experimental paradigm was achieved by using a parallel port to send different trigger pulses for each event from Matlab to the EEG amplifiers.

To control the signal quality, a visual inspection of the EEG data was done both continuously during the measurement as well as after the experiment in Brainstorm (Tadel et al., 2011). We searched for EMG artifacts by examining time course and topography of single-trial data. Channels with strong contamination with EMG artifacts were excluded from further analysis in single subjects; on average, we rejected 1.03 ± 1.75 (mean ± SD) channels.

During signal processing, the EEG data and markers were down-sampled from 5 kHz to 1 kHz (time-frequency analysis) or 500 Hz (voltage plots). Channels were re-referenced to their common average. The signal was filtered with a Butterworth high-pass filter of fourth order with a cut-off frequency of 0.5 Hz.

The indices of the correct and false responses were extracted and aligned on the response EMG. The time point of the EMG onset was found by applying a threshold based retrospective search on the arm EMG channels.

### Intracranial EEG recording, localization and preprocessing

Recording of intracranial EEG signal was done either with Compumedics amplifiers (Singen, Germany) at the epilepsy center in Freiburg, Germany (2 kHz sampling rate), or with Schwarzer Epas amplifiers (Munich, Germany) and Nicolet EEG C-series amplifiers (Pleasanton, USA) at the epilepsy center of the Motol University Hospital in Prague, Czech Republic (512 Hz sampling rate). The depth electrodes used for recording had platinum-iridium contacts (DIXI Medical, Lyon, France & AD-TECH, Racine, WI, USA).

The preprocessing was done as for the noninvasive data, with the difference that the channels were re-referenced bipolarly between the respective neighbors.

The stereotactic depth electrodes were localized with the help of their post-implantation MRI or CT artifacts in a normalized and co-registered MRI of each patient as described in Pistohl et al. (2012). After transformation to the MNI coordinate system, cytoarchitectonic probabilistic maps were calculated with the SPM anatomy toolbox (Eickhoff et al., 2005, 2006, 2007) to assign the electrodes to specific brain regions. This method accounts for inter-subject differences in brain anatomy and allows the comparison on a group level with high precision (Amunts et al., 2007).

Electrodes positioned inside a seizure onset zone or showing frequent interictal activity, as identified by experienced epileptologists, were excluded from further analysis. Of 885 bipolar referenced channels from 9 patients, we removed 76 channels because they could not be assigned to a specific brain region, 53 channels which were positioned inside a seizure onset zone, 59 channels because of frequent interictal activity, and 7 channels due to technical problems or position outside the brain. Approximately 20 % of the remaining electrodes were identified as lying within white matter. As a control, we also did the analyses of the intracranial data after splitting white and gray matter electrodes; however, as we found that white matter electrodes, especially those near boundary areas, were able to sample a number of significant effects, and further did not change any findings, we did not exclude these channels from further analysis. Thus, 690 sites were available for further analysis.

The visualization of intracranial and scalp EEG electrodes in relation to the cortex surface was done with Brainstorm (Tadel et al., 2011). The intracranial electrodes were visualized on an ICBM152 brain template (Mazziotta et al., 2001; Fonov et al., 2009).

### Time-frequency analysis

Time-resolved spectral power was computed with a multitaper method (Thomson, 1982) with a window length of 500 ms, a step size of 50 ms and two Slepian taper functions. Trial averages were computed with a median function, which proved to be a robust method in past EEG studies (Ball et al., 2008). To visualize error-related power modulations we baselined all time-frequency bins in each error trial by the corresponding bin of the median correct response trial (i.e., by dividing the error responses by median of the correct responses). Thus, the motor-related activity common to both response types canceled out. For calculation of error-related activity or power, this method of baselining was applied.

### Exclusion and statistical matching of ocular artifacts

Ocular movements were recorded with the EyeLink 1000 plus (SR Research Ltd., Ottawa, Canada, RRID:SCR_009602), enabling binocular eye movement recordings with a high-speed infrared illuminator and camera; the sampling rate for binocular recording was 500 Hz.

For the extraction of microsaccades, an algorithm as described by Engbert and Kliegl (2003) was used with the default minimal microsaccade duration threshold of 12 ms. Time points of blinks and saccades were identified with the algorithms provided by the EyeLink recording software.

All trials in which a blink occurred within the time window of 0.5 s prior and 2 s after the response were excluded from further analysis to ensure that the EEG signal in the analyzed time frame was not contaminated with blink artifacts. On average, 112 ± 164 (mean ± SD) trials were rejected across subjects.

To exclude possible influences of saccades and microsaccades on high-frequency EEG correlates, we matched each time bin of the correct and error trials after the time frequency decomposition. This was done by incrementally removing time bins from the correct condition, which always had more trials. At each iteration step, the time bin which contributed the most to the residual difference, as calculated by a sum of squares of saccade and microsaccade counts, was removed. This process was repeated until the p-value calculated with a sign test comparing error and correct condition was greater than 0.3.

### Statistical analysis

For visualization of median voltage, the standard error of the median was calculated per condition using bootstrap sampling (Moore and McCabe, 1989) with 1000 re-samples, using the 2.5^th^ and 97.5^th^ percentile of the population as standard error. Median time or frequency bins which had a significant difference between the correct and erroneous condition were identified with the Wilcoxon rank sum test (Mann and Whitney, 1947) in single subjects or with a two-sided sign test (Dixon and Mood, 1946) across subjects. Significance of median power changes within individual conditions was computed with a two-sided sign test. Whenever multiple comparisons were done, we estimated the positive false discovery rate (pFDR) for each p-value (Benjamini and Hochberg, 1995; Storey, 2002, 2003).

## Results

Here we show for the first-time error-related high-gamma responses in a noninvasive EEG study (with 31 healthy subjects) and additionally compare and corroborate them with intracranial EEG measurements (in 9 patients with pharmacoresistant epilepsy).

### Error-related voltage and spectral power modulations in noninvasive EEG

In the following, we show an overview of the dynamics and topography of voltage and spectral power modulations, as averaged over the 31 healthy subjects included in this study, in correct responses (Fig. 2), erroneous responses (Fig. 3) as well as the difference between the two conditions, i.e., error-related activity (Fig. 4). The spectral power responses are shown up to 300 Hz in the time interval from 0.7 s before until 1.5 s after the response. For these and all comparable topographical results, the positive false discovery rate was calculated across 128 channels, 24 frequency bands, and 61 time points (1 s before response onset until 2 s after response onset in steps of 50 ms). Error-related HGB activity had its maximum approximately between 60 and 90 Hz (Fig. 2-4, yellow background), or more narrowly located between 70 and 80 Hz (Fig. 2-4, blue background). Based on these observations, we decided to use these frequency ranges for a closer inspection of HGB responses in the following analyses

**Figure 2:**
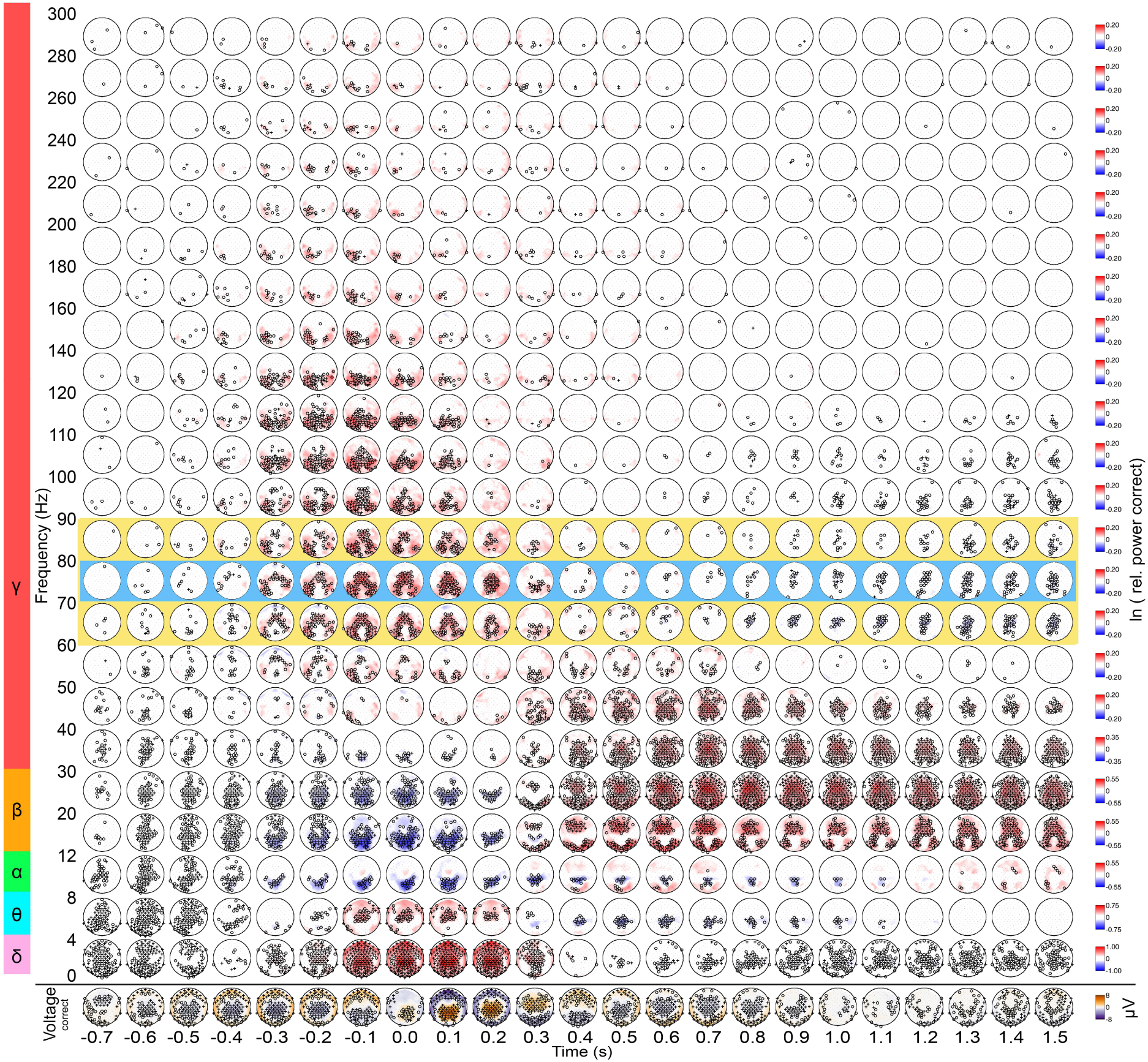
Topography of voltage and spectral power modulations in a correct response. The circular plots represent a top view of the head with the nose pointing up. At the bottom, the average response is plotted (median of 31 subjects). Above, the median spectral power modulations relative to the time from -1 s to -0.5 s prior to the response, are depicted for 23 frequency bands from 0 to 300 Hz. Red color indicates higher power than in the baseline, blue color indicates lower power than in the baseline period. Electrodes with significant power modulations are marked with “o” or “+” (sign test, pFDR<0.05 or pFDR<0.01, respectively). The data is aligned to response onset.

**Figure 3:**
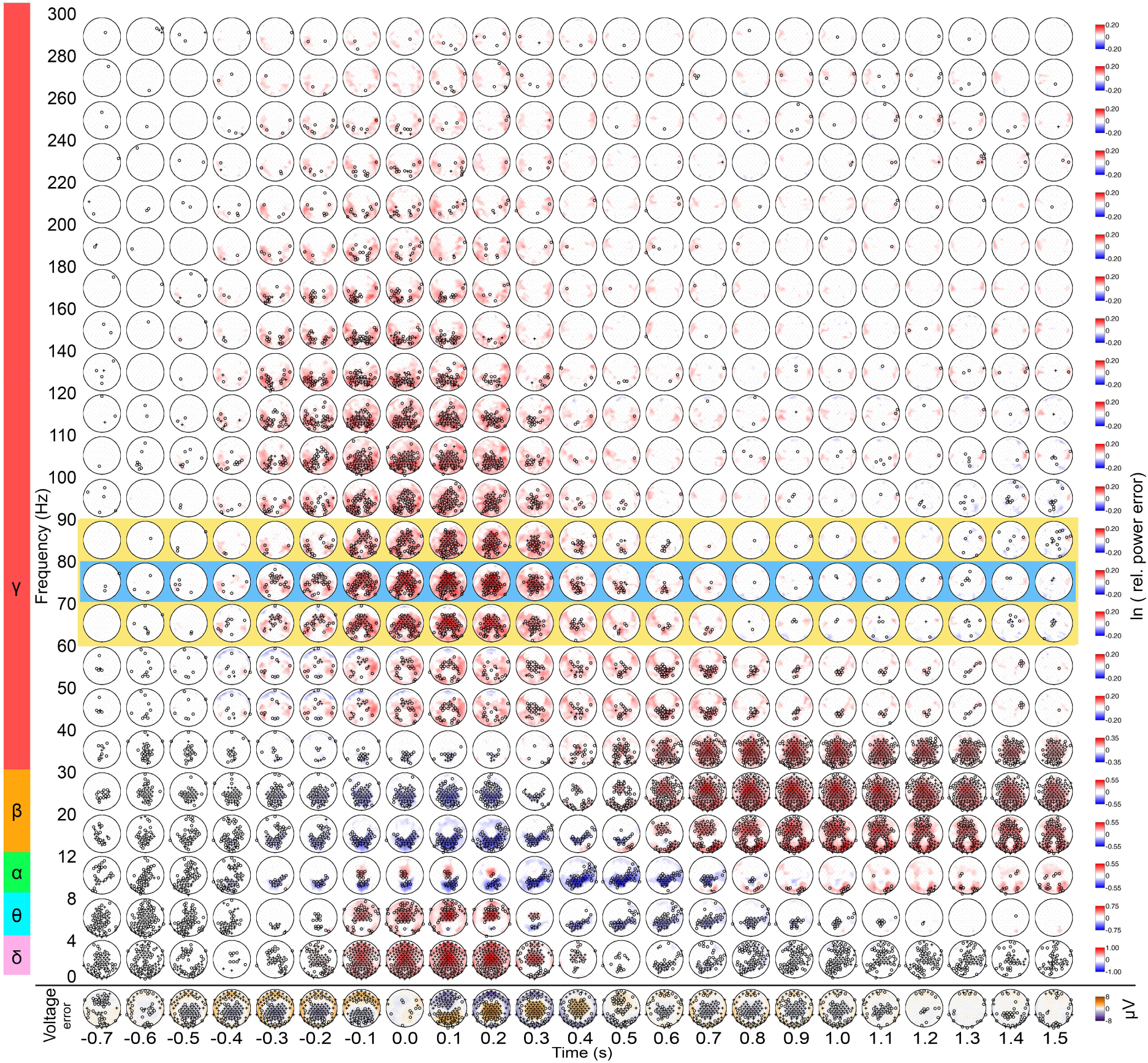
Topography of voltage and spectral power modulations in an error response. All conventions as in Figure 2.

**Figure 4:**
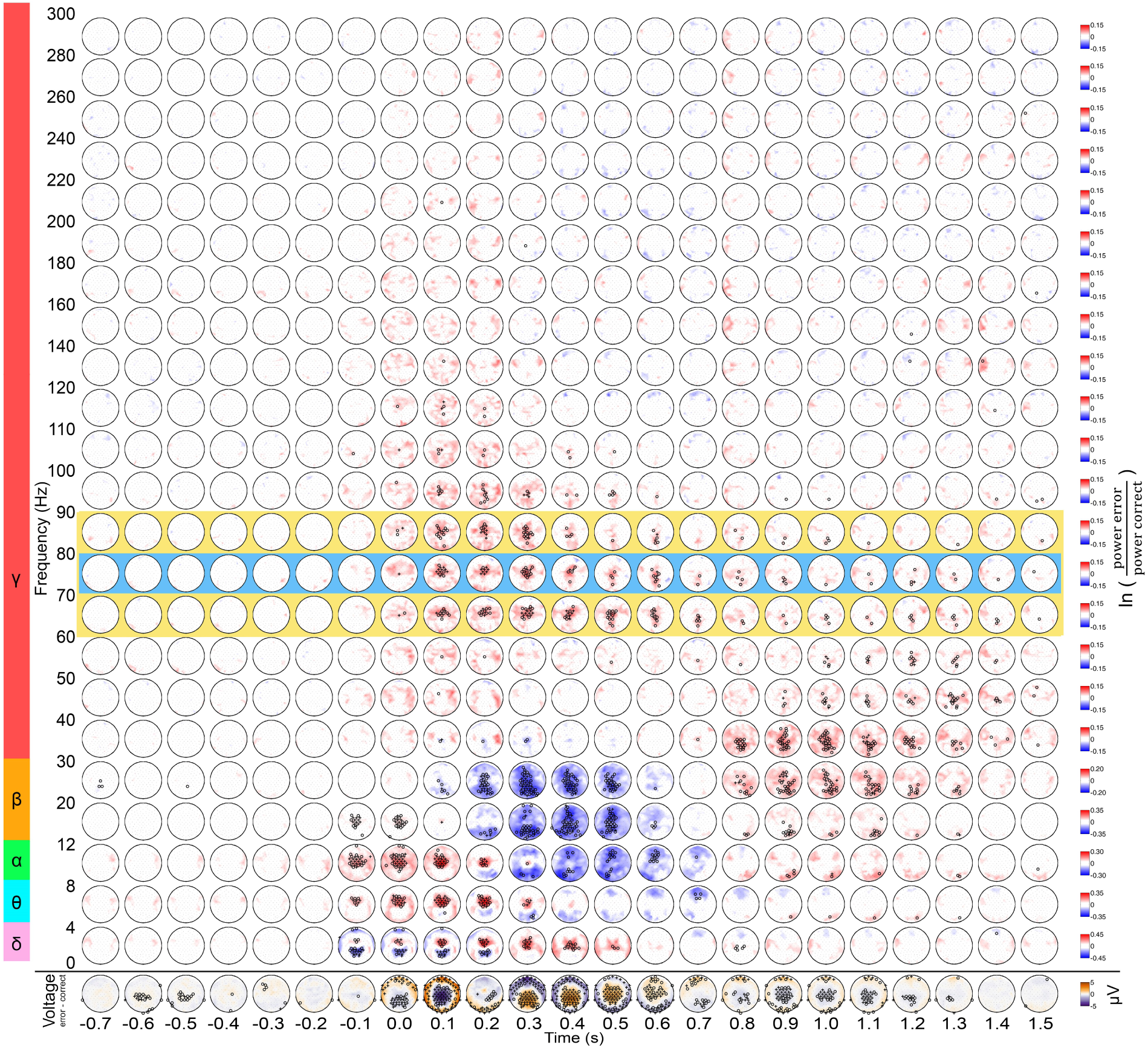
Topography of error-related voltage and spectral power modulations. Conventions as in Figure 2. At the bottom, voltage differences between error and correct response are shown topographically (median of 31 subjects). Above, median error-related spectral power modulations are shown, i.e., each time-frequency point of the error response was baselined with the same point of the median correct response. Red color indicates higher power in error responses, while blue color indicates higher power in correct responses. Electrodes with significant differences between error and correct condition are marked with “o” or “+” (sign test, pFDR<0.05 or pFDR<0.01, respectively). The data is aligned to response onset.

Comparing error and correct conditions, significant differences became evident. While there was lower power at parietal channels in the delta band (< 4 Hz) during and after errors, fronto-central channels exhibited an error-related power increase in the delta and the theta band (4-8 Hz). In the alpha band (8-12 Hz), a spatially more widespread power increase during and shortly after the erroneous response occurred, followed by an attenuation with lower power compared to correct responses. Within the beta band (12-30 Hz), a widespread power decrease concurred with the Pe. In the low-gamma band (LGB, 30-50 Hz), differences were mostly apparent at a later time, starting 800 ms post-response in the form of a power increase in the 30-40 Hz frequency range. Simultaneously with the late LGB increase, a second (with the ERN/Ne being the first) significant negative voltage deflection was seen at central channels.

Crucial to the present study, in the high-gamma band (HGB, > 50 Hz), a significant error-related power increase occurred at fronto-central channels shortly after the response onset, and shifted to more central and parietal areas over the course of a few hundred milliseconds, where the error-vs-correct relative HGB power stayed significantly increased until up to 1.5 s after the response. Similar high-gamma increases were observed in errors both after incongruent and congruent stimuli classes (data not shown). After an initial HGB power increase, there was a significant midline power decrease in correct responses after 800 ms until at least 1.5 s (Fig. 2). This power decrease was not observed after erroneous response (Fig. 3)

Fig. 5 depicts the median frequency profile of the correct, error and error/correct conditions for a central region of interest (ROI).

**Figure 5:**
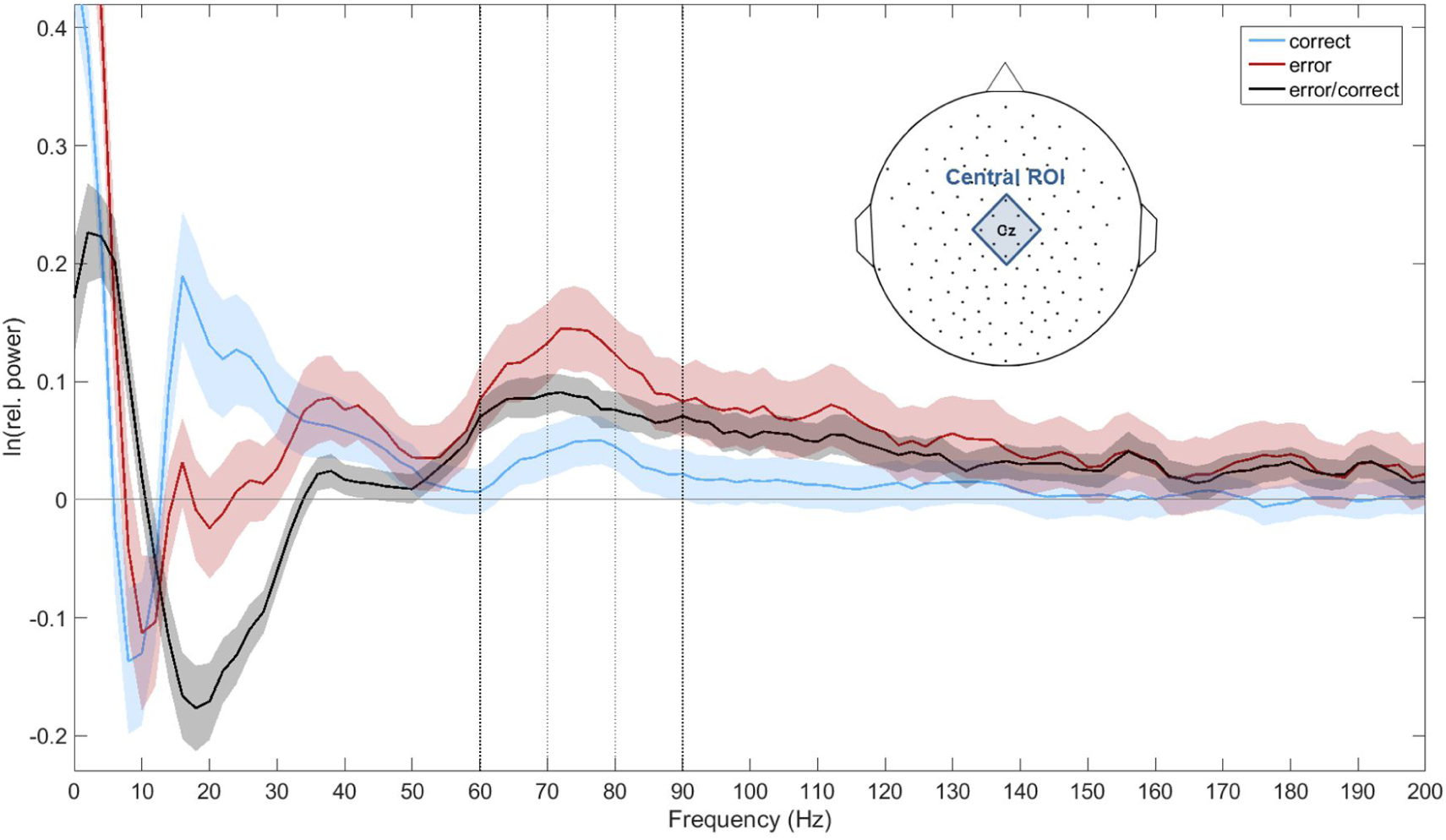
Frequency profile of spectral power modulations in an Eriksen flanker task. The average frequency profile for the correct (blue), error (red) and error/correct condition (black) is shown averaged over a central ROI (FCz, FCC1h, FCC2h, C1, Cz, C2, CCP1h, CCP2h, CPz) during a time of interest between 0 s and 0.6 s relative to response onset. The SEM is plotted semitransparent. While relative power changes were highest in the delta (0-4 Hz) and theta (4-8 Hz) range, strong modulations were also observed in the alpha (8-12 Hz), beta (12-30 Hz) and high-gamma (> 50 Hz) range. The two frequency ranges of interest in the high-gamma range are marked with bold (60 to 90 Hz) and thin (70 to 80 Hz) dashed lines, respectively.

Error-related voltage and spectral power modulations of selected frequency bands in surface EEG are shown in Fig. 6. Topography and time course of the classical ERN/Ne and Pe components can be seen in the upper part. In the lower part of the figure, the relative power differences between error and correct condition are plotted in five frequency ranges of interest at the time point of the ERN/Ne and Pe components.

**Figure 6:**
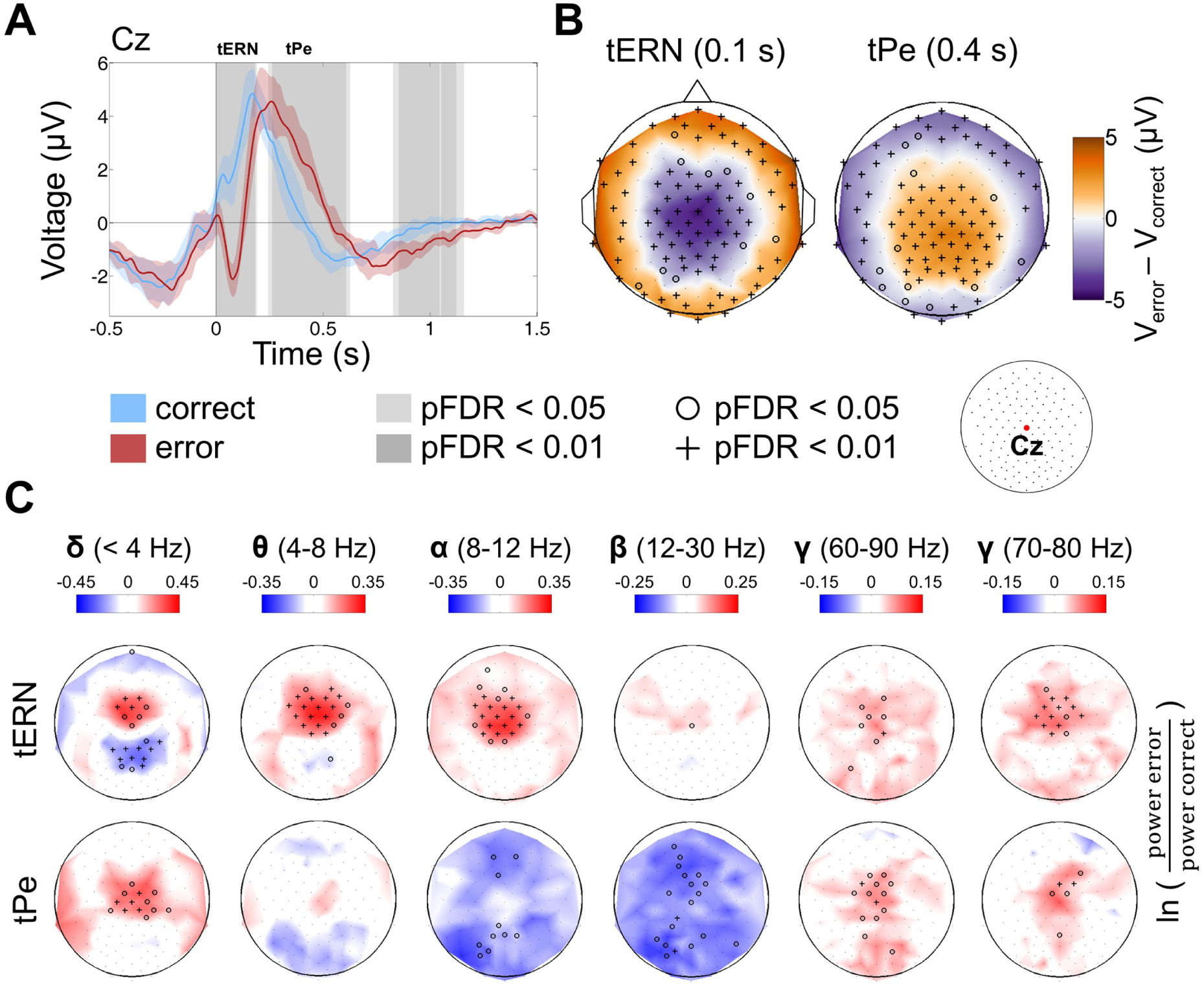
Error-related voltage and spectral power modulations in high-density EEG. **A)** Single channel plot (electrode position Cz) locked to response EMG onset. The median correct response is shown in blue, the median erroneous response in red. The standard error of the median is plotted semitransparent. The vertical dotted line marks the EMG onset. Times with significant differences between the error and correct condition are shown with a gray background; pFDR values were calculated across 1501 time points (-1 s to 2 s, 500 Hz). **B)** Topographical map of the event-related potentials at the time of the ERN/Ne (t_ERN_) and the Pe (t_Pe_) components. Median voltage difference between error and correct trials is shown color coded. Electrodes with significant differences are marked with “o” or “+” (sign test, pFDR<0.05 or pFDR<0.01, respectively); pFDR values were calculated across 31 time points (-1 s to 2 s in steps of 100 ms) and 128 channels. **C)** Error-related spectral power modulations in the theta, alpha, beta, and high-gamma range. Logarithmic relative power of the median erroneous response baselined with median correct response is shown color coded. Significant sites as determined (sign test) are marked as in (B); pFDR values were calculated as in Fig. 3. Error-related effects in the high-gamma range had a maximum at fronto-central sites at the time of the ERN/Ne peak; at the time of the Pe maximum, the HGB increase was shifted to midline electrodes in a parietal direction.

The high-gamma band response, shown here for the time window between 70 and 80 Hz, exhibited significantly higher power in error than correct responses. In the time window of the ERN/Ne maximum, the error-related HGB power was centered at fronto-central channels, while it started to shift to more posterior areas around the maximum of the Pe.

We calculated Spearman's rank correlation level between the ERN/Ne and Pe amplitude and the error-related spectral modulations in the 70-80 Hz band across subjects. The calculation was done on the signals of the channels with the maximal amplitudes of the components of interest, which was FCz for the ERN/Ne and CPz for the Pe component. There was no significant correlation found. Spearman’s rho for the ERN/Ne with error-related HGB components was r_s_ = -0.05 and p = 0.78, and for Pe r_s_ = 0.10 and p = 0.58.

### Effects of eye movements on high-gamma signals

During the experiments, a great number of miniature eye movements were recorded for each subject. After discarding those events near blinks, on average 6300 ± 2500 (mean ± SD) microsaccades were recorded per subject. To examine the spectral power modulations related to this class of eye movements more closely, we also examined EEG data aligned to the onset of the microsaccades. Fig. 7 shows the group median of those trials. Both at fronto-polar and parieto-central EEG channels, there was a broadband gamma increase during microsaccade (and saccade, data not shown) onset. This increase was prominent from 25 to 120 Hz. In lower frequency bands, e.g., in the delta band, we observed a power decrease, which continued until 500 to 600 ms after the microsaccade onset.

**Figure 7:**
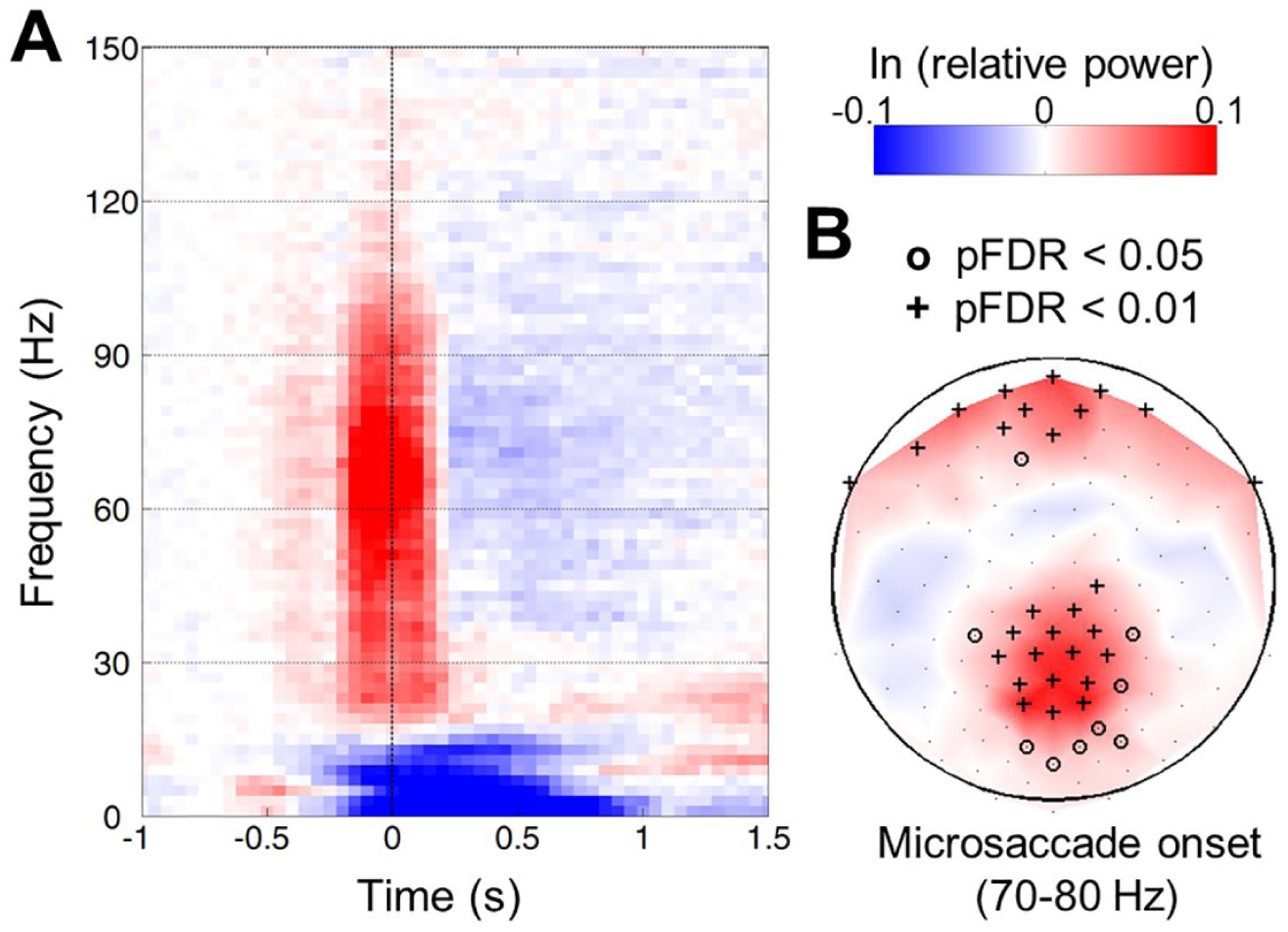
Microsaccade-related time-frequency spectrum in scalp EEG. The plots show a group median of the 31 subjects; the data is aligned to microsaccade onset. **A)** Time-frequency plot of a parieto-central channel (Pz). The log-power (color-coded) was taken relative to a baseline from -1 s to –0.5 s. **B)** Topography of 70-80 Hz high-gamma relative power at the time of microsaccade onset. Electrodes with significant differences are marked with “o” or “+” (sign test, pFDR<0.05 and pFDR<0.01, respectively); pFDR values were calculated as in Fig. 3.

Next, in each subject we matched the correct and error condition to have the same amount of microsaccades and saccades within each time-frequency bin of the EEG data after the multitaper analysis. To examine the effect of this measure on the results, we compared the error-related high-gamma activity in matched and unmatched data (Fig. 8).

**Figure 8:**
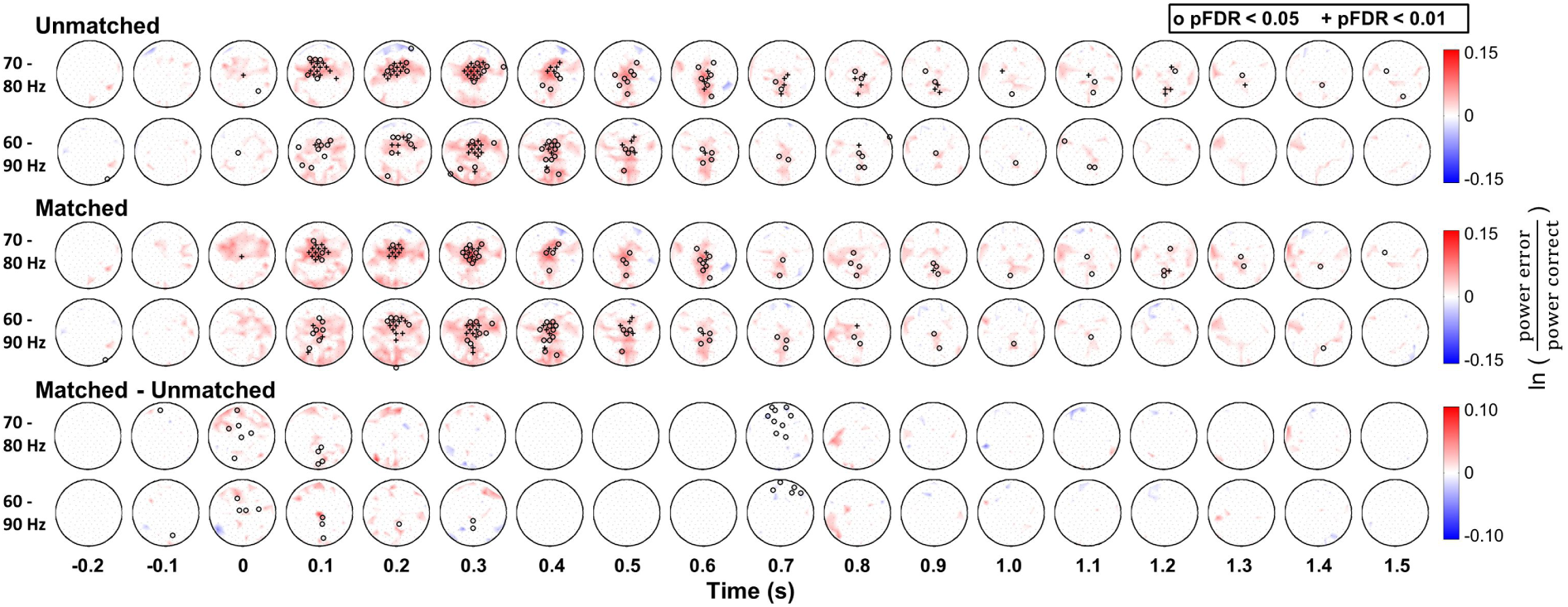
Influence of saccade and microsaccade matching on error-related high-gamma in noninvasive EEG. All plots show group results of the 31 subjects, aligned to EMG response onset. In the top row, error-related high-gamma activity in the 70-80 Hz and 60-90 Hz range is plotted for the data before matching. The middle row depicts the same results after matching each time bin for microsaccade and saccade frequency. In the bottom row, the differences between unmatched and matched data are shown. Significant sites are marked as in Fig. 4.

After the matching, the distribution of electrodes with significant changes in the high-gamma band was only minimally altered, and the overall pattern remained the same. There were only very few significant differences between the original and the matched data. We conclude the error-related effects cannot be attributed to the occurrence of saccades and microsaccades.

### Error-related high-gamma activity: frontal vs. parietal regions

We further wished to compare error-related high-gamma activity at frontal and parietal regions as the two major foci of the HGB response. To this aim, we averaged HGB activity in a frontal and a parietal region of interest (ROI).

Error-related HGB power in the frontal ROI reached its maximum around 150 to 200 ms after response EMG response onset. The same frequency band in the parietal ROI showed a later peak at 600 to 700 ms. The HGB power in the parietal ROI was significantly increased until up to 1.3 s after the EMG response onset. In the time around the EMG onset, error-related HGB activity in the frontal ROI was significantly (sign test, p<0.01) stronger than in parietal regions.

### Confirmation in Intracranial EEG Measurements

Intracranial EEG measurements in 9 patients yielded a multitude of error-related changes. For comparability, we concentrated the analysis of error-related high-gamma activity in intracranial EEG to the 60-90 Hz band, as the frequency of interest in noninvasive EEG. An overview of the electrode locations of all 9 patients and sites with significant changes in this range is shown in Fig. 10 together with examples of single-channel time-frequency responses.

**Figure 10:**
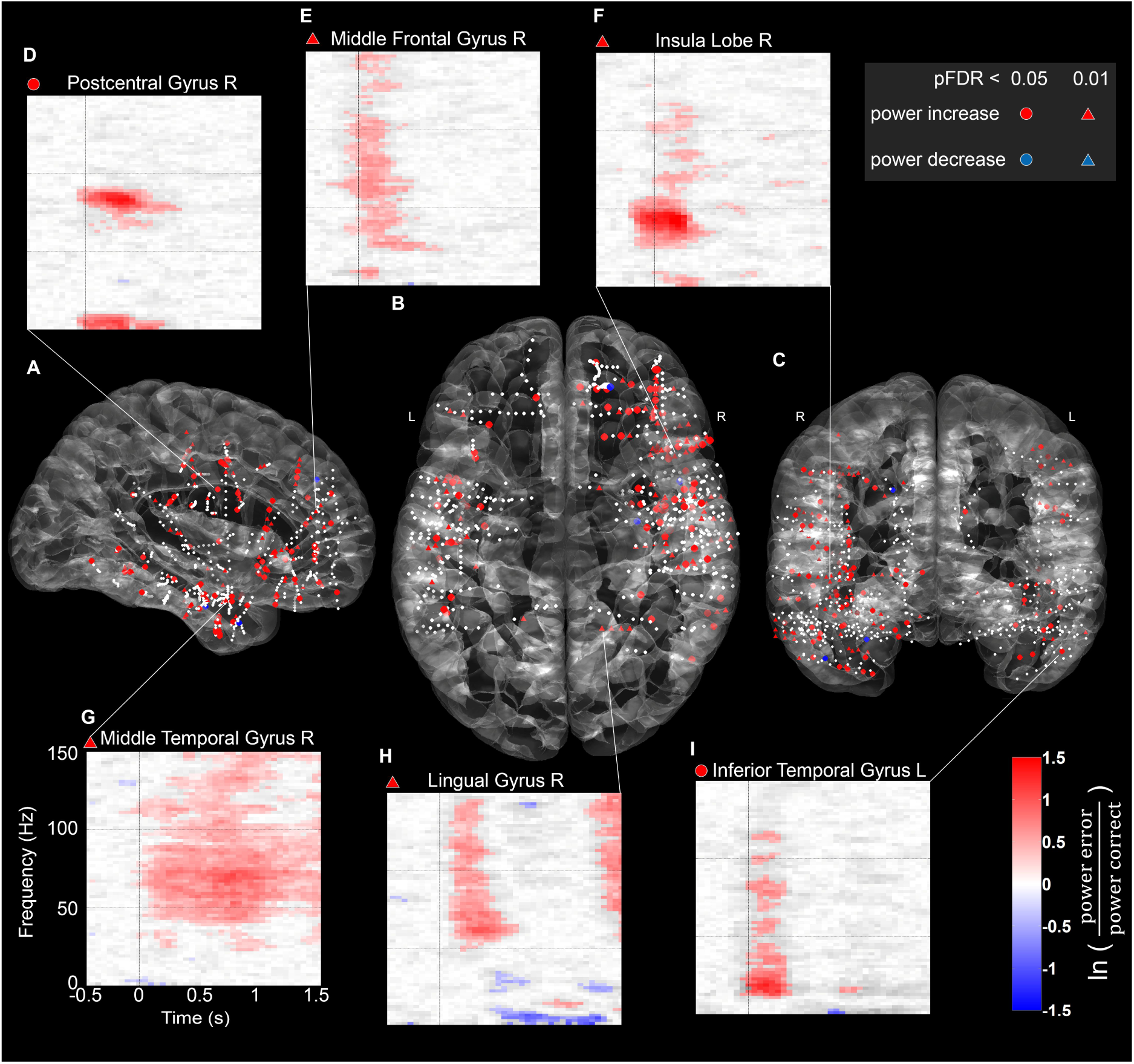
Distribution of error-related high-gamma (60-90 Hz) responses in intracranial EEG. Depth electrodes of 9 epilepsy patients are plotted in white on the semi-transparent ICBM152 brain template from sagittal (A), axial (B), and coronal (C) views. Sites with significant activations in the 60-90 Hz range in the time of interest between 0 s and 0.6 s after an error, as identified in noninvasive EEG, were marked with colored circles or triangles (sign test, pFDR<0.05 or pFDR<0.01, calculation of pFDR values across all channels). The marker color indicates whether there was a relative power increase (red) or decrease (blue) compared to correct responses. Exemplary single-channel time-frequency plots are shown above and below together with their assignment to anatomical areas; time-frequency bins with pFDR>0.05 are shown grayed out (calculation of pFDR across 51 times (-0.75 s to 1.75 s, 50 ms steps) and 126 frequencies (0-250 Hz, 2 Hz steps). Color scale as in Fig. 6 C.

Error-related high-gamma modulations were found in multiple areas across the cortex, mostly manifesting as power increases relative to correct responses. As is noticeable in the exemplary time-frequency plots, we observed different types of error-related HGB modulations in intracranial EEG. Firstly, responses with a clear maximum which was restricted to a small area in time and frequency, as the examples from the postcentral gyrus and the insula (Fig. 10 D,F). Secondly, responses with a transient, broadband gamma increase, e.g., in the middle frontal gyrus, the lingual gyrus or the inferior temporal gyrus (Fig. 10 E,H,I). Thirdly, responses with a broad-band gamma increase and with a long duration of around 2 s, e.g., in the middle temporal gyrus (Fig. 10 G).

More significant HGB modulations were located in the right hemisphere; however, it is to note that our sample consisted of more electrodes in the right hemisphere (443 sites after bipolar re-referencing & channel rejection) compared to the left hemisphere (247 sites). Across the 9 patients, we observed an average error rate of 21.09 ± 0.11 (mean ± SD) % with the left hand, and 20.81 ± 0.10 % with the right hand; there was no significant difference with p=0.46 (paired one-sided t-test).

To get an overview of the significance of single-channel observations, we calculated the ratio of significant responses within 19 regions of interest (Table 1).

**Table 1:** **Significant error-related 60-90 Hz power modulations 489 in different areas**. For 19 ROIs, the percentages of channels with significant (pFDR<0.05) 60-90 Hz HGB error-related power modulations, averaged across the time window from 0 s to 0.6 s after the response (FDR-corrected overall channels, as in Fig. 10), are listed together with the total number of channels (n) included per area.

| Area | n | % positive | % negative |
| --- | --- | --- | --- |
| Heschls gyrus | 2 | 100.00 | 0.00 |
| Lingual gyrus | 8 | 62.50 | 0.00 |
| Middle frontal gyrus | 35 | 45.71 | 0.00 |
| Precentral gyrus | 43 | 44.19 | 0.00 |
| Insular cortex | 47 | 38.30 | 0.00 |
| Inferior frontal gyrus | 71 | 33.80 | 0.00 |
| Postcentral gyrus | 40 | 30.00 | 0.00 |
| Fusiform gyrus | 39 | 28.21 | 0.00 |
| Middle cingulate cortex | 4 | 25.00 | 0.00 |
| Supramarginal gyrus | 25 | 24.00 | 0.00 |
| Inferior temporal gyrus | 72 | 23.61 | 0.00 |
| Hippocampal area | 60 | 18.33 | 1.67 |
| Orbitofrontal cortex | 73 | 17.81 | 0.00 |
| Superior temporal gyrus | 35 | 17.14 | 0.00 |
| Superior frontal gyrus | 22 | 13.64 | 4.55 |
| Middle temporal gyrus | 117 | 12.82 | 0.85 |
| Anterior cingulate cortex | 24 | 8.33 | 0.00 |
| Rolandic operculum | 25 | 4.00 | 0.00 |
| Precuneus | 6 | 0.00 | 0.00 |

For 19 ROIs, the percentages of channels with significant (pFDR<0.05) 60-90 Hz HGB error-related power modulations, averaged across the time window from 0 s to 0.6 s after the response (FDR-corrected over all channels, as in Fig. 10), are listed together with the total number of channels (n) included per area.

Error-related high-gamma power increases were observed in 18 of the 19 ROIs, with a range of 4 % to 100 % of involved electrodes in the different areas. Power decreases were only observed in 3 regions, ranging from 0.85 % to 4.55 % of electrodes.

We further compared error-related spectral power modulations of nearby intracranial EEG and scalp EEG electrodes in the range of the motor cortex (Fig. 11).

**Figure 11:**
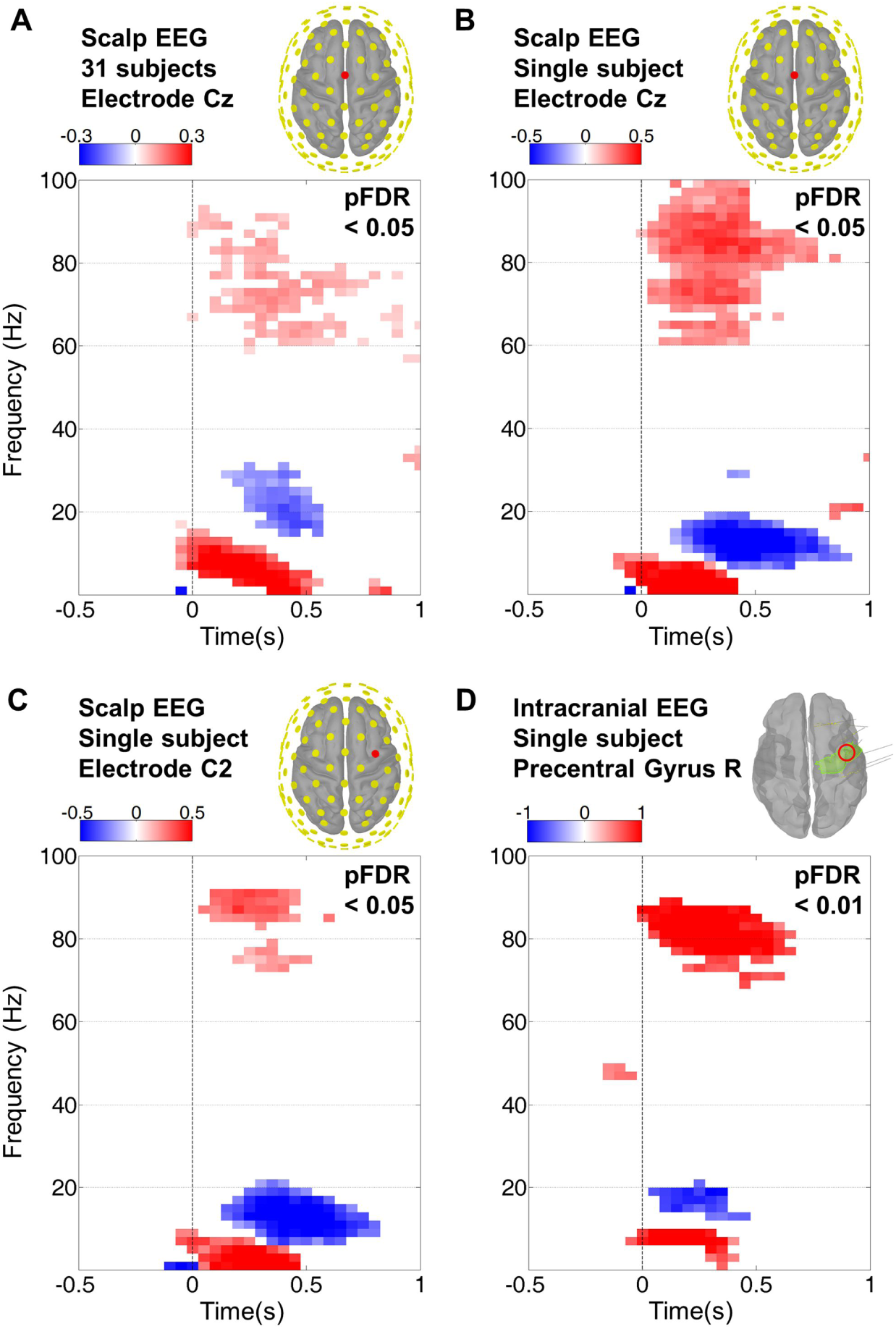
Comparison of intracranial EEG with scalp EEG in group and single subject results. The color scale depicts the logarithmic relative power of the median error-related response baselined with the correct response. Only significant power modulations are shown with the threshold as labeled in the top right corner of the respective plot. Calculation of pFDR-values was done with an FDR-corrected Wilcoxon rank sum test (single subjects) and an FDR-corrected sign test (group results). **A)** Group median (31 subjects) of error-related spectral power modulations at electrode Cz. **B)** Single-subject error-related spectral power modulations in EEG at electrode Cz. **C)** Single-subject error-related spectral power modulations in EEG at electrode C2. **D)** Single-subject error-related spectral power modulations in intracranial EEG within the right precentral gyrus in close proximity of the C2 electrode standard position.

Both in scalp EEG at central positions and intracranial EEG in the precentral gyrus, significant error-related power modulations were observed in the delta, theta, alpha, beta, and high-gamma band. In these examples, the spectral patterns of nearby intracranial and noninvasive EEG channels were very similar at both group and singlesubject level (Fig. 11). In intracranial EEG, the high-gamma activity had the strongest error-related power modulation, while in noninvasive measurements, the lower frequency bands showed a greater difference between correct and error responses.

### Fine-grained dynamics of error-related activity in intracranial EEG

For a closer look at the spatio-temporal progression of error-related activity across the brain, we analyzed the time course of error-related low- and high-spectral power modulations in the intracranially recorded data of all 9 patients (Fig. 12).

**Figure 12:**
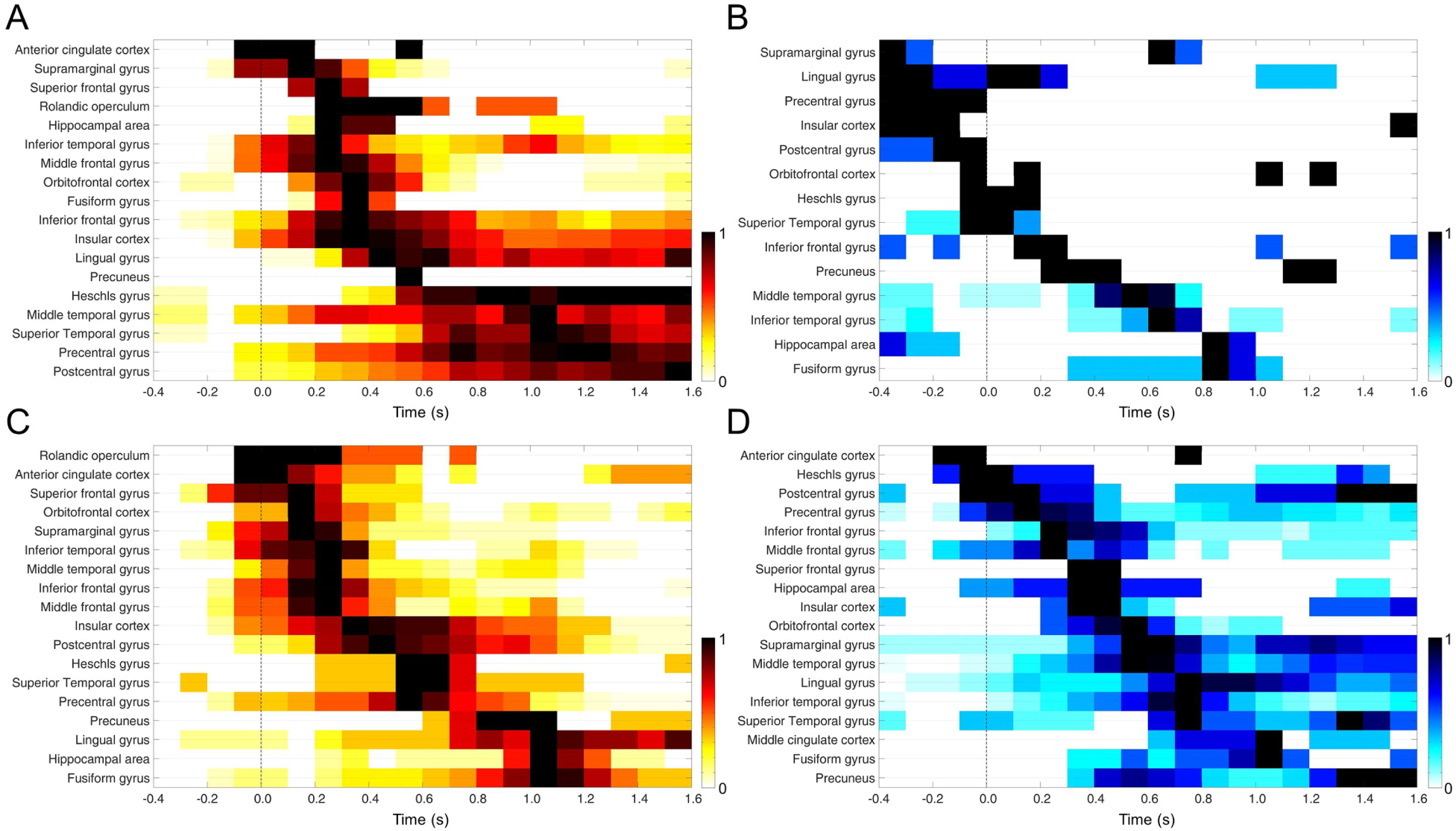
Time course of significant error-related power modulations in intracranial EEG. For each area, the total number of significant increases (red) and decreases (blue) in all patients were normalized; the areas were sorted according to the time with the maximal count (equal to 1) in ascending order. Time course of HGB power modulations in the range 50-120 Hz are shown at the top (A, B), low-frequency components below 30 Hz are shown at the bottom (C, D). The data is aligned to response onset.

It is apparent that low and high frequencies, as well as power increases and decreases, differed in their spatial distribution over time. Subareas in frontal, temporal, and parietal brain regions were activated at various time points before, during and after the error response. Error-related increases in the high-gamma range (Fig. 12 A) especially exhibited a temporal development that started at frontal surface und deep areas, including the ACC, and then advanced to both parietal and temporal regions of the brain. Notably, increased error-related HGB activity in hippocampal areas peaked 0.3 s after the response, while decreased hippocampal HGB activity (Fig. 12 B) occurred -0.4 to -0.2 s before the response and again 0.8 to 1.0 s after the response. Overall, intracranial EEG portrays a consistent but much more complex picture of error processing compared to noninvasive data.

## Discussion

Converging evidence indicates an important role of HGB activity in cortical function (Baçar et al., 2001; Herrmann et al., 2004a; Fries, 2005; Canolty et al., 2006; Crone et al., 2006; Jensen et al., 2007; Jerbi et al., 2009b; Buzsáki and Wang, 2012). Studies on human HGB activity were initially restricted to intracranial EEG (Szurhaj and Derambure, 2006; Miller et al., 2007; Whitham et al., 2008). A growing body of literature, however, indicates that HGB activity can be measured noninvasively using EEG (Muthukumaraswamy, 2013). HGB activity has been measured noninvasively during sensory stimulation (Cobb and Dawson, 1960; Heinrich and Bach, 2004; Scheller et al., 2005), cognitive processing (Friese et al., 2013; Long et al., 2014), executed (Ball et al., 2008; Darvas et al., 2010; Nottage et al., 2013) and imagined motor tasks (Smith et al., 2014). This wealth of noninvasive HGB demonstrations underscores that it is in principle possible to measure HGB activity noninvasively. While error-related HGB activity has been demonstrated in intracranial EEG (Milekovic et al., 2013; Bastin et al., 2017), here we show error-related HGB activity noninvasively.

In the present study, we examined error processing in the human brain as reflected in HGB activity, based on measurements using noninvasive and intracranial EEG. In both, we found significant error-related modulations of high-gamma power. Noninvasive 128-channel EEG in an electromagnetically shielded cabin enabled us to reveal the global topography and dynamics of event-related potentials and spectral power modulations, while we used intracranial EEG to validate the noninvasive findings, and additionally to probe local fine-grained activity patterns.

### Error-related low-frequency responses in noninvasive EEG

Our findings generally reproduced the spectral power modulations in the delta, theta, alpha and beta bands in noninvasive EEG as reported by previous studies (Luu et al., 2004; Yordanova et al., 2004; Kolev et al., 2005; Trujillo and Allen, 2007; Koelewijn et al., 2008; Carp and Compton, 2009). Further, our results also revealed several unreported topographical features.

For example, we observed an error-related delta band pattern characterized by the co-occurrence of increased and decreased power in anterior and posterior regions, respectively (Fig. 4,6). In the high-beta and low-gamma band (20 - 40 Hz), we observed an error-related power increase in a late time window, starting 800 ms after the response (Fig. 4,6). Interestingly, around 1000 ms after response onset, a second (with the ERN/Ne being the first) smaller but significant (p<0.01) negative deflection in the error-related potential occurred at midline EEG channels (Fig. 4, bottom row) which thus might be termed “Ne1000”. The maximal low-gamma band response occurred at roughly the same time with a focus on midline channels. Together, these examples illustrate that an optimized EEG procedure applied to a suitably large group of subjects can reveal a range of additional significant features of the error-related response that may be useful to consider in future studies.

### Error-related high-frequency responses in noninvasive EEG

Both in group results and single subjects, we found significant error-related HGB power increases. Although high-gamma activity was seen in both correct and erroneous trials, it was significantly stronger in the erroneous trials. The error-related high-gamma response presented itself as an early fronto-central power increase, followed by a shift to parieto-central areas, where the HGB power after errors was significantly larger than than after correct responses over an extended period (up to 1.5 s, see Fig. 4,9). Importantly, the spatio-temporal dynamics differed clearly from those of theta, alpha, beta, and low-gamma responses, and high-gamma response were not correlated to ERN/Ne and Pe amplitudes across subjects, pointing to a unique functional role.

We controlled thoroughly for ocular artifacts in our noninvasive EEG experiments. Firstly, by using high-resolution binocular eye tracking, we were able to exactly match each time bin of the error-related and non-error-related time-frequency-resolved data for both microsaccade and saccade frequency and thus avoid a differential effect of ocular potentials (Yuval-Greenberg et al., 2008) on the error-related high-gamma signals. Secondly, the topography of microsaccade-related HGB effects showed a qualitatively different spatial pattern compared to the error-related HGB responses (Fig. 7, 8). Thus, we conclude that our error-related HGB responses cannot be explained as ocular artifacts.

Further, we also think that it is highly unlikely that our error-related HGB responses reflect EMG contamination. First, the error-related HGB power did not exhibit the rather flat, broadband power increase typical of EMG contamination (Goncharova et al., 2003), but rather a strong maximum between 60 and 90 Hz. Second, the spatial distribution of the high-gamma increase with the maximum over the midline differed from that to be expected for EMG, which has its emphasis on peripheral electrodes close to the muscles (Goncharova et al., 2003; Whitham et al., 2007).

### Error-related high-gamma responses in intracranial EEG

Error-related intracranial high-gamma activity was found at multiple locations in the brain, showing much larger amplitudes than the extracranial counterparts. These areas overlapped strongly with areas where intracranial error-related potentials and HGB increases were previously reported (Brázdil et al., 2005; Bastin et al., 2017). In addition to gamma increases, we also observed responses with a decreased relative high-gamma power in relation to errors. They often followed a HGB power increase, possibly representing a systematic suppression aftereffect. As saccadic artifacts in intracranial EEG are mainly limited to the temporal pole (Jerbi et al., 2009a), our intracranially recorded error-related HGB increases together with the results reported in 6 patients by Bastin et al. (2017) clearly show the existence of such responses on the cortex. Furthermore, as illustrated by Fig. 12, intracranial and scalp EEG channels placed above central brain regions showed coinciding time-frequency patterns in their error-related responses (see Fig. 12). Together, these observations lend additional support to the validity of our noninvasive data. One limitation of such a comparison is, of course, that not all areas can be observed in noninvasive EEG recordings. While we assume signals from the hippocampus and insular cortex to be virtually undetectable in noninvasive EEG, signals from cingular areas might be recordable from the surface (Ball et al., 1999). Generally, the superficial regions of the cortex areas can be expected to have a greater influence on the scalp EEG than subcortical areas (Nunez et al., 1997).

The time course of intracranial activations (Fig. 13) confirmed that frontal regions were activated rather early, while parietal (as well as temporal) areas became active later, in line with a downstream role in error processing. Intracranial data also clearly showed that, expectedly, cortical error processing is more complex than what can be observed in noninvasive EEG. Hippocampal high-gamma was significantly decreased prior to the actual error, and increased after the error. This could signify an interaction of error and memory systems, consistent with a role of gamma in memory functions (Howard et al., 2003; Jensen et al., 2007; Sederberg et al., 2007; Kucewicz et al., 2014, 2017). Furthermore, Fig. 13 also demonstrates that lower HGB power in pre-, postcentral, and supramarginal gyri as well as the insular cortex may precede errors. One explanation for that could be that high-gamma power could indicate a pre-activation or “readiness” of a brain area.

### Interpretation of high-gamma band activity

Broadband power modulations in intracranial EEG are positively correlated with the local single-neuron firing rate in humans (Manning et al., 2009) and can be explained by current models (Bédard et al., 2006; Ray et al., 2008; Miller et al., 2009), which assume a linear low-pass kernel to quantify the influence of single synaptic events on the population signal. Insofar as the underlying spiking activity can be assumed to sufficiently irregular (Softky and Koch, 1993; London et al., 2010), the presynaptic action potentials can be modeled as a Poisson process, which has a broad spectrum contributing equally at all frequencies. Some characteristics of the average kernel proposed in these previous works, such as the cutoff frequency and the order of the low-pass filtering, have been experimentally investigated (Miller et al., 2009). Our present EEG results are consistent with broadband changes across a wide frequency range (Fig. 2-5), possibly reflecting similar mechanisms as previously described in intracranial data.

In addition to broadband changes, inhibitory coupling may lead to gamma oscillations manifesting in more narrow-banded peaks in different ways. If fluctuations of the membrane potential are small, e.g., if the neurons are driven tonically, inhibitory connections can serve to synchronize the activity in the population, leading to regular and globally synchronous firing (Wang and Buzsáki, 1996; White et al., 1998; Kopell et al., 2000).

In contrast, the mechanism of a collective oscillatory instability relies on inhibitory feedback arriving with a temporal delay *d*; oscillatory frequencies are in the range of 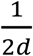 and 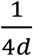 (Brunel and Hakim, 1999; Tiesinga and José, 2000), thus well typically in the high gamma range. The oscillation in these states persists on the population level -it is manifest in the pairwise correlations between neurons - while single cells exhibit irregular firing (Brunel and Hakim, 1999; Brunel, 2000; Tiesinga and José, 2000; Brunel and Wang, 2003).

Another network mechanism that can give rise to gamma oscillations relies on the feedback loop between excitatory (E) and inhibitory (I) neurons (Wilson and Cowan, 1972), requiring a strong coupling from E to I and from I to E (reviewed in Buzsáki and Wang, 2012). The resulting oscillations in spiking network models are typically in the lower gamma range (Brunel and Wang, 2003). This mechanism has also been employed in a non-linear rate model to explain the dependence of gamma oscillation frequency on the stimulus size in visual cortex, exploiting the modulation of the intra-cortical E-I feedback loop through the neuronal non-linearity (Kang et al., 2010). Notably, the gamma frequency peak around 80 Hz observed in our data shows a similarity to the gamma frequency profiles predicted by the model by Brunel (2000, Fig. 9 A).

**Figure 9:**
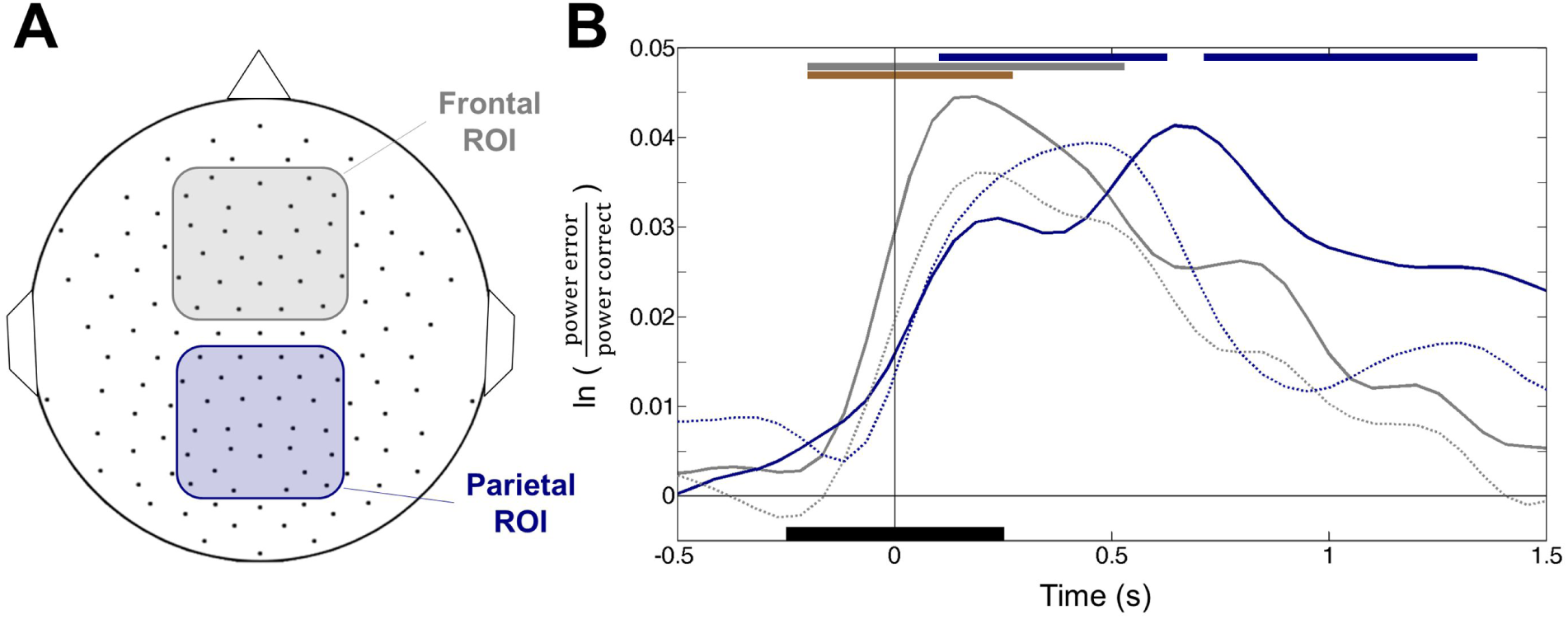
Time course of error-related 70-80 Hz HGB activity in frontal and parietal regions. **A)** Illustration of the frontal region of interest (ROI) including 25 electrodes (gray) and a parietal ROI with 27 electrodes (blue). **B)** The median error-related 70-80 Hz high-gamma power is plotted for both ROIs in the respective colors; error-related 60-90 Hz power is plotted with dashed lines in the same colors. The bars above indicate times when the 70-80 Hz HGB activity in the frontal (gray), parietal (blue) or difference of both ROIs (brown) was significant (sign test, pFDR<0.01, calculated as in Fig. 6 A). Time point zero represents the response EMG onset. The black bar around 0 s at the bottom of the plot indicates the temporal width of the Gaussian window function (0.5 s) used for the time-resolved spectral density estimation during moving average-calculation, explaining the smoothness of the curves above.

Also, more realistic networks comprised of several layers (Potjans and Diesmann, 2014) are able to generate an oscillation in the same frequency range: A mean-field analysis shows that the network mechanism is here a subcircuit comprised of excitatory and inhibitory neurons in layers 2/3 and 4 (Bos et al., 2016). The mathematical analysis of these networks also exposes why the power of the oscillation is strongly influenced by the tonic drive to the network, predominantly to layer 4. An error-related signal changing the tonic drive to the local oscillation-generating network may thus be a candidate mechanism behind the observed modulations in the power spectra. Such a mechanism would predict a co-modulation of the multi-unit firing rate with the increase of gamma power, as previously empirically observed (Nir et al. 2007) and very much in line with our observation of a co-occurrence of both, broad band changes and additionally enhanced gamma power around 80 Hz (Fig 2-5). Thus, a distinction between a broadband increase of gamma power by the firing rate alone from a modulation of intrinsic oscillatory properties of the circuit may thus not be as clear cut.

From our present results we still cannot draw firm conclusions about the mechanisms giving rise to the enhanced gamma activity that we have prominently observed around 80 Hz and beyond. To gain such insight in future studies, our noninvasive EEG method may prove to be a useful tool to study gamma responses and they mechanistic underpinnings.

### Functional relevance of error-related HGB activity

Our findings reveal sustained, significant HGB activity over the parietal cortex that substantially outlasted the duration of the Pe, up to at least 1.5 s after response onset. This raises the question whether this specific timing may hint at a possible functional role of this late high-gamma response during error processing. Generally, processing of behavioral errors involves several prominent functional sub-processes. These range from precursors of error detection, such as the evaluation of actual and intended action outcomes, including perceptual evidence accumulation, over the explicit error detection itself, to the evaluation of error importance, error-related learning, and behavioral adjustment (Holroyd and Coles, 2002; Carbonnell and Falkenstein, 2006; Taylor et al., 2007). Much research so far has focused on how these processes relate to the ERN/Ne and Pe complex. Recent evidence, based on systematic reward manipulations that targeted the criterion which participants used to decide whether or not to report errors, has linked the Pe to the strength of accumulated error-related evidence (Steinhauser and Yeung, 2010). Late, sustained parietal high-gamma activity could thus reflect processes downstream to error evidence accumulations, such as behavioral adjustment and motor learning. This speculation could be tested in future studies utilizing EEG-based HGB mapping as described in our present study.

### Conclusion & Outlook

The fact that error-related HGB signals are detectable with noninvasive EEG opens up a much wider avenue of research than would be feasible with intracranial recordings alone, particularly in healthy subjects and also in populations of patients with different neuropsychiatric disorders, such as social phobias, autism spectrum disorders, schizophrenia, depression, as well as obsessive-compulsive disorder – all of which have been connected with alterations in the neural response to behavioral errors (Alain et al., 2002; Hajcak and Simons, 2002; Hajcak et al., 2003; Henderson et al., 2006; Ruchsow et al., 2006; Shiels and Hawk, 2010; Riesel et al., 2011). Implantation of intracranial electrodes is confined to a much smaller group of patients undergoing pre-neurosurgical evaluation, in most cases for the treatment of focal pharmacoresistant epilepsy. Our findings however clearly highlight the unique value of these intracranial recordings. For example, they allow assessment of brain structures that are difficult or even impossible to probe electrophysiological noninvasively, such as insular cortex and the hippocampal formation.

Our findings could help understanding the mechanisms behind human error processing. Further, they could also help in the decoding of errors from single-trial EEG using machine learning algorithms to improve the performance of brain-machine interfacing (Ferrez and Millan, 2008; Kreilinger et al., 2012; Spüler et al., 2012; Milekovic et al., 2013; Völker et al., 2017). To dissect underlying mechanisms, an examination of phase-amplitude coupling between high-gamma power and lower frequency bands (Canolty et al., 2006; Cohen et al., 2008; Tort et al., 2010; Friese et al., 2013) during error processing could provide valuable information. More generally, we suggest that parallel investigations of error processing with both noninvasive as well as invasive recordings, and with attention to both classical error-related potentials as well as high-frequency signatures of error processing, might prove as the most fruitful way towards understanding the neural basis of this fundamental facet of cognition.

## Acknowledgements

This work was supported by DFG grant EXC1086 BrainLinks-BrainTools, Baden-Württemberg Stiftung grant BMI-Bot, Graduate School of Robotics in Freiburg, Germany and the State Graduate Funding Program of Baden-Württemberg, Germany.

